# Benchmark for simple and complex genome inversions

**DOI:** 10.1101/2025.11.28.691176

**Authors:** Siyuan Cheng, Fritz J Sedlazeck

## Abstract

**Background:** Inversions represent a consequential yet under-characterized form of structural variation, with roles in genomic disorders, evolution, and genome instability. However, their detection remains technically challenging, particularly within repeat-dense regions and for complex multi-breakpoint events. A lack of dedicated, high-quality benchmarks has hindered algorithmic improvement, performance comparison, and robust biological interpretation.

**Results:** Here, we present a comprehensive, multi-genome benchmark for simple and complex inversions derived from Strand-seq and phased long-read based assemblies across five reference samples, with breakpoint refinement using haplotype-resolved long-read assemblies. This tiered resource spans a broad spectrum of inversion classes, sizes, and zygosity states, and captures challenging genomic contexts including segmental duplications, inverted repeats, and composite rearrangements. We used this benchmark to systematically assess leading structural variant callers and alignment strategies across short-read, PacBio HiFi, and Oxford Nanopore data. Performance varied substantially by inversion class and genomic context: simple inversions were recovered with high sensitivity at sufficient coverage, whereas complex and heterozygous events remained difficult. Sniffles2 and Severus achieved the strongest recall for complex inversions, despite increased false-positive rates. We additionally benchmarked two commonly used long-read alignment pipelines (Minimap2 and VACmap), demonstrating that the mapper choice has a substantial impact on inversion detection in repetitive regions.

**Conclusion:** Together, this work provides the first unified, high-resolution inversion benchmark and reveals clear strengths and limitations of current methods across platforms. Our resource establishes a foundation for principled tool development, evaluation, and tuning, enabling the community to more accurately resolve inversion variation and its biological and clinical consequences.

## Introduction

Inversions are a class of chromosomal structural variants (SVs) in which a continuous segment of DNA is reversed end-to-end relative to the reference orientation[1–4]. Inversions are relatively rare in the human genome, yet they can exert substantial biological and clinical effects[5]. Well-known cases include pathogenic inversions in IDS (Hunter Syndrome)[6], the DUP-TRP/INV-DUP rearrangements in MECP2 Duplication Syndrome[2,7], the F8 inversion leading to Hemophilia A[8,9], and the inversion in acute myeloid leukemia leading to gene fusion[10,11]. Inversions can also contribute to disease susceptibility in more subtle ways, which manifest as developmental and neurological disorders in carriers’ offspring[12,13].

Several molecular mechanisms can lead to the formation of inversions. A common mechanism is non-allelic homologous recombination (NAHR), in which recombination occurs between two sequences that share high similarity but are located at different positions and inverted orientations in the genome[4]. NAHR between inverted repeats or segmental duplications (SDs) can flip the intervening segment, producing a recurrent inversion[4]. Inversions can also form via non-homologous end joining (NHEJ) or retrotransposon activity[4,14,15]. Complex replication-based mechanisms, such as fork stalling and template switching (FoSTeS) or microhomology-mediated break-induced replication (MMBIR), can generate more complicated inversion structures, involving multiple breakpoints or co-occurrence with other structural variants[4,16].

Despite their importance and impact across multiple diseases, their discovery is limited by the need for multi-technology and methodology approaches[2,16,17]. This is because inversions are often located around complex or repetitive genomic regions of the genome, which facilitate their emergence[16]. Traditional short-read DNA sequencing (SRS) technology struggles with the large or complex inversions that are often of great interest[1,18]. The emerging long-read sequencing (LRS) technologies show improvements in the detection of inversion breakpoints with longer read length[1,2]. However, it still struggles with the most intractable repetitive structures when an inversion is flanked by repeats longer than even the long reads, where assembly, or mapping, can fail[16]. Strand-seq has proven useful for detecting inversions, but is limited by its low resolution of breakpoints and high false-positive rate in short inversions (<5Kb)[16,19,20]. Besides technology-based challenges, analysis remains surprisingly ignorant of inversions. For example, common aligners often are not optimized for inversions, and variant calling methods often rely blindly on the aligner itself[1,21]. This is also because of a lack of benchmarks targeted for inversions. Current SV benchmark efforts, like Genome in a Bottle (GIAB)[22,23] and Human Genome Structure Variant Consortium (HGSVC)[24,25], have been focusing on insertion and deletion of SV or SNVs exclusively. This lack of benchmarks hinders method development and comparison of methods. Thus not challenge the field enough to develop new approaches to cope with inversions.

In this paper, we introduce an inversion benchmark across 5 samples (HG002, HG02818, HG03486, HG00733, NA19240) that facilitates the uptake of inversion detection and thus hopefully the broader study of inversions and complex rearrangements. For this, we utilize Strand-seq[20] with breakpoint refinement facilitated by long-read assemblies[26] to build a robust benchmark resource for simple and complex inversions. We classified inversions into three confidence tiers and stratified them by genomic context, size, genotype, and repeat content. Utilizing the benchmark set, we systematically evaluated current inversion callers across calling strategies, aligners, and sequencing technologies. Our results highlight critical gaps in inversion detection: recall for inversions in large repeats (>5 kb) remains low, heterozygous inversions are markedly harder to detect compared to homozygous ones, and the choice of aligners heavily impacts the precision and recall, etc. Overall, this benchmark provides an urgently needed foundation for improving inversion detection methodologies.

## Results

### A Tiered High-Confidence Inversion Set Through Assembly-Based Refinement of Strand-seq Loci

To construct an accurate benchmark for inversions, we utilized Strand-Seq data information across five samples (HG002, HG00733, HG02818, HG03486, NA19240) to identify the loci where an inversion occurs[20]. Furthermore, we refined the exact breakpoints by aligning the corresponding HPRC[26,27] phased assemblies against the GRCh38 reference genome and developed a custom script to assess breakpoints (**Methods**). The process involves breakpoint identification using split-read and soft-clip evidence of the aligned phased assemblies, genotyping of paternal and maternal haplotype calls, and systematic filtering and annotation, culminating in a hierarchically merged final callset (**Figure 1A, Methods**).

**Figure 1:**
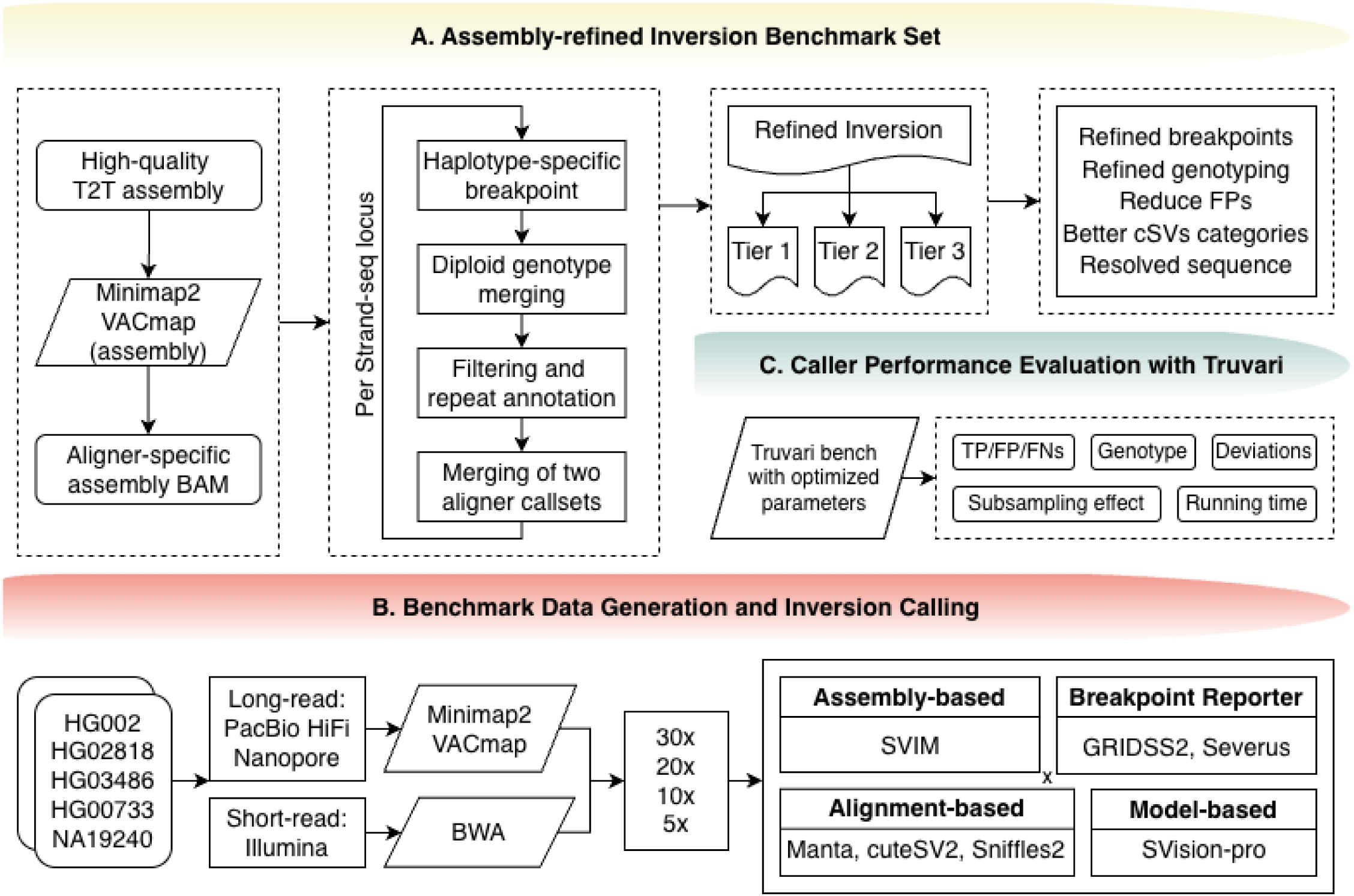
Overview of the inversion benchmarking pipeline. **(A) Assembly-refined inversion benchmark set construction.** High-quality T2T assemblies were mapped back to the reference (Minimap2/VACmap), yielding assembly-specific BAMs. Within each Strand-seq locus, haplotype-aware breakpoint detection and diploid genotype merging were performed, followed by filtering, repeat annotation, and merging of calls from two aligners to produce a tiered (Tier 1–3) refined inversion VCF. **(B) Benchmark data generation and inversion calling.** We analyzed five human samples sequenced with PacBio HiFi, Oxford Nanopore, and Illumina platforms at different coverages. Long reads were aligned with Minimap2 or VACmap, and short reads with BWA, to produce aligner-specific BAMs. Inversion callers comprised assembly-based methods, alignment-based tools, and a model-based caller. **(C) Caller performance evaluation with Truvari.** Each caller’s output was benchmarked against the refined VCF using Truvari bench with inversion-optimized parameters, generating precision, recall, F1 score, genotype concordance, breakpoint, and length shift statistics, TP/FP/FN counts. Subsampling analyses assessed the impact of coverage.

The resulting dataset was stratified into three confidence tiers to facilitate downstream benchmarking and analysis. Tier 1 represents the most confident set, comprising inversions with breakpoints highly concordant with Strand-seq and the assembly data. Tier 2 includes inversions with fewer breakpoint concordances and overall generally higher complexity. Tier 3 consists of loci with a Strand-seq signal but no supporting evidence in the assembly-based alignments from either Minimap2[28] or VACmap[21]. Tier 3 calls were thus excluded from this benchmark set.

Overall, we processed 125-154 inversion candidates per sample across five samples (**Table 1 and Supplementary Table 1-2**). Each sample had 62-92 Tier 1 inversions, which mainly contained simple inversion events with and without repeats flanking them. These are predominantly smaller inversions, with most events falling within the 50 bp to 100 kb range, where median values range from 3.9 Kb to 6.1 Kb, and maximum values range from 304.5 Kb to 584.9 Kb (**Table 2**). Tier 2 contained 53-82 inversions and complex rearrangement events. In contrast to Tier 1, Tier 2 is enriched for larger inversions, with most events in the 100 kb to >1 Mb size bins where median values range from 90.3 Kb to 189.6 Kb, and maximum values range from 4.49 Mb to 6.26 Mb. Furthermore, Tier 2 inversions are mostly heterozygous and are more often flanked by large repeats (>5 kb), underscoring the challenges in resolving and genotyping complex variants in repetitive genomic regions.

**Table 1:**
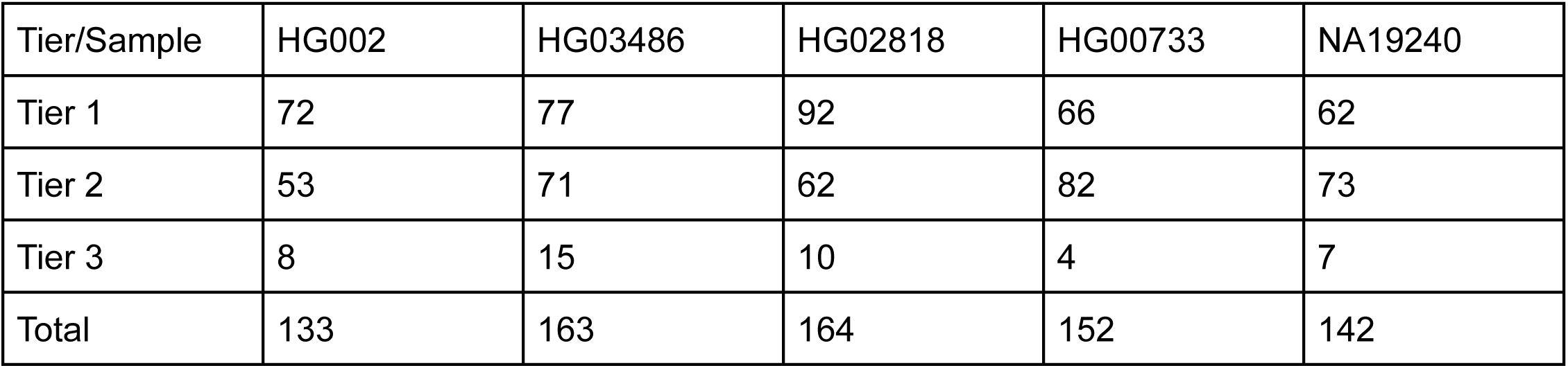
Assembly-refined inversion numbers across samples.

**Table 2:** Stratified assembly-refined inversion set for HG002 and HG03486.

| Sample | Tier | Total | Length |  |  |  | Genotype |  | Repeat |  |
| --- | --- | --- | --- | --- | --- | --- | --- | --- | --- | --- |
|  |  |  | 50bp<br>-10<br>kb | 10-100<br>kb | 100kb-1Mb | >1Mb | Hom | Het | Repeat<br>≥5Kb | Repeat<br><5Kb |
| HG002 | Tier 1 | 72 | 45 | 15 | 12 | 0 | 32 | 40 | 35 | 37 |
|  | Tier 2 | 53 | 0 | 20 | 24 | 9 | 7 | 46 | 34 | 19 |
|  | Tier 3 | 8 | 0 | 0 | 0 | 0 | 0 | 0 | 0 | 0 |
| HG03486 | Tier 1 | 77 | 49 | 25 | 3 | 0 | 32 | 45 | 30 | 47 |
|  | Tier 2 | 71 | 3 | 24 | 32 | 12 | 4 | 67 | 50 | 21 |
|  | Tier 3 | 15 | 0 | 0 | 0 | 0 | 0 | 0 | 0 | 0 |

We conducted further manual inspections using IGV[29] based on miminap2 and VACmap alignments of the assemblies and the raw reads. This showed overall good concordance and alignment support on breakpoint resolution, genotyping accuracy, correcting false positives, and better capture of complex inversions. For instance, our pipeline consistently refined breakpoints to base-pair resolution, as seen at chr2:138247429 (**Figure 2A**), and improved genotyping accuracy for complex events, such as chr17:45578419 (**Figure 2B**). Nevertheless, limitations persist for the most complex Tier 2 inversions, particularly those embedded within centromeres or flanked by extensive SDs due to the incomplete sequence on the GRCh38 genome or splits in the assemblies themselves.

**Figure 2:**
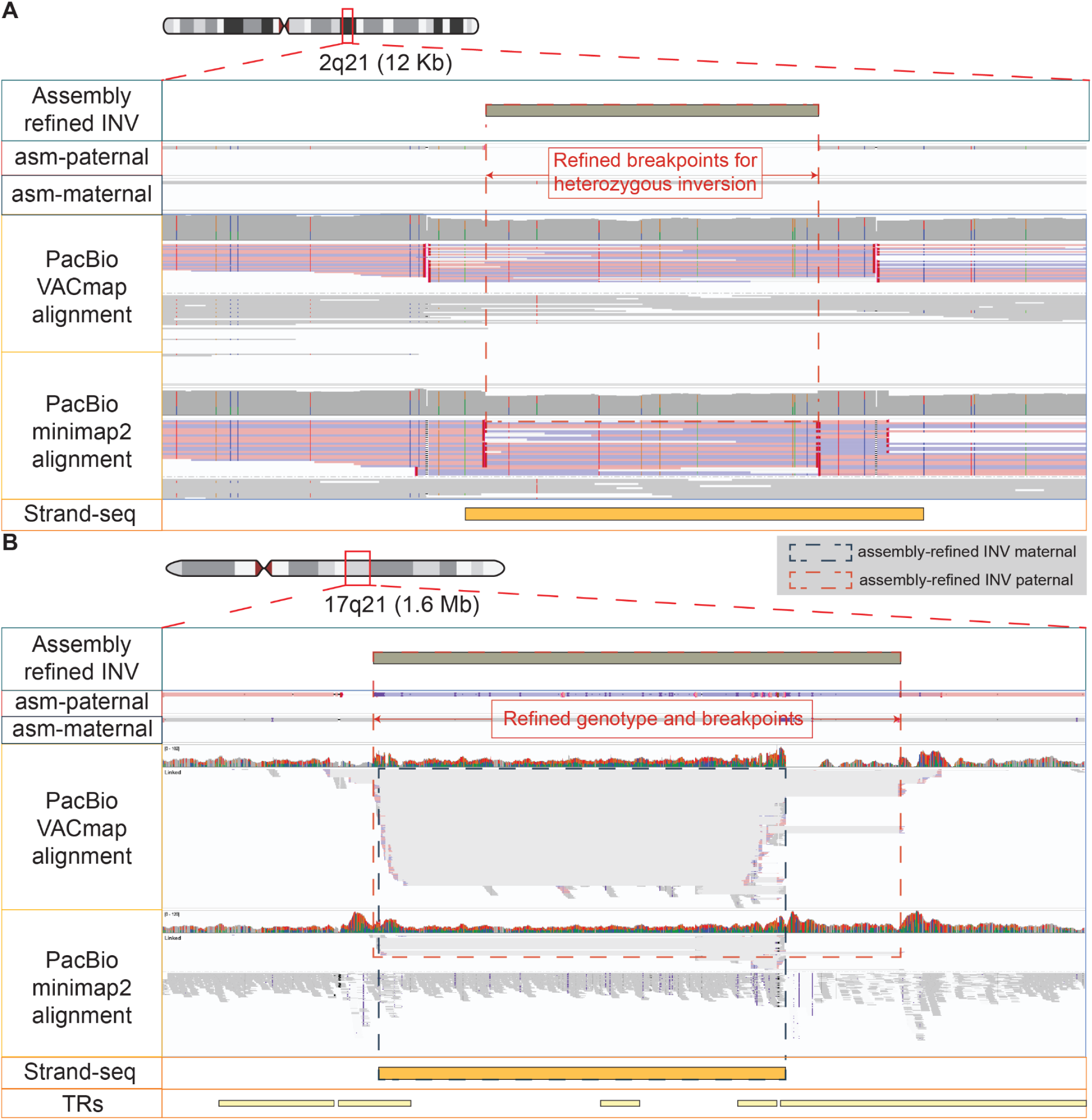
Assembly-based refinement improves breakpoint resolution and genotyping of Strand-seq inversion calls. **(A) A 3.6 Kb heterozygous inversion on chromosome 2 (2q21).** The assembly-based pipeline refines the initial Strand-seq call, defining the breakpoints to base-pair resolution. Tracks show the final assembly-refined inversion, the consistent paternal and maternal haplotype assemblies (asm-paternal, asm-maternal), supporting PacBio read alignments (VACmap and minimap2) where colored reads (blue and pink) are linked reads indicating inverted orientation, and the original Strand-seq locus. **(B) A complex 900 Kb heterozygous inversion on chromosome 17 (17q21.31).** The assembly-based approach accurately corrects the genotypes and refines the breakpoints of this complex event. The divergence between the paternal and maternal assemblies, supported by PacBio read alignments, clarifies the structure and heterozygous state of the inversion, improving upon the initial Strand-seq call.

### Benchmarking strategies and overall results

We investigated inversion detection across seven SV calling methods, including two short-read callers (GRIDSS2[30], Manta[31]), and five long-read callers (cuteSV2[32], Severus[33], Sniffles2[2], SVIM[34], SVision-pro[35]), representing different calling strategies (**Figure 1B**). Sequencing data for Illumina, PacBio, and Oxford Nanopore (ONT) across five human samples (HG002, HG00733, HG02818, HG03486, NA19240) were downloaded and subsampled at four different coverages (30x, 20x, 10x, 5x). We evaluated the performance of Minimap2[36] and VACmap[21] across the seven SV callers using Truvari[37] (Methods).

We discuss the results in terms of recall (ie, correctly identified inversions), precision (ie, TP proportion among calls), and overall F1 score (ie, harmonic mean of the precision and recall) (**Methods**). We adjusted the allowable reference distance (refdist) for small (50 bp–10 kb), medium (10 kb–100 kb), and large (>100 kb) inversions, given that many inversions are flanked by repeats and thus their accurate breakpoints are often elusive. This refined approach requires stricter criteria for smaller variants, where high resolution is expected, while accommodating the inherent ambiguity of breakpoints in larger structural changes (**Methods**).

To start evaluating inversion detection, we benchmarked 30x whole-genome datasets by examining recall (**Figure 3**), focusing on HG002 and HG03486 and their stratifications, where other samples’ results can be found in **Supplementary Figure 1**.

**Figure 3:**
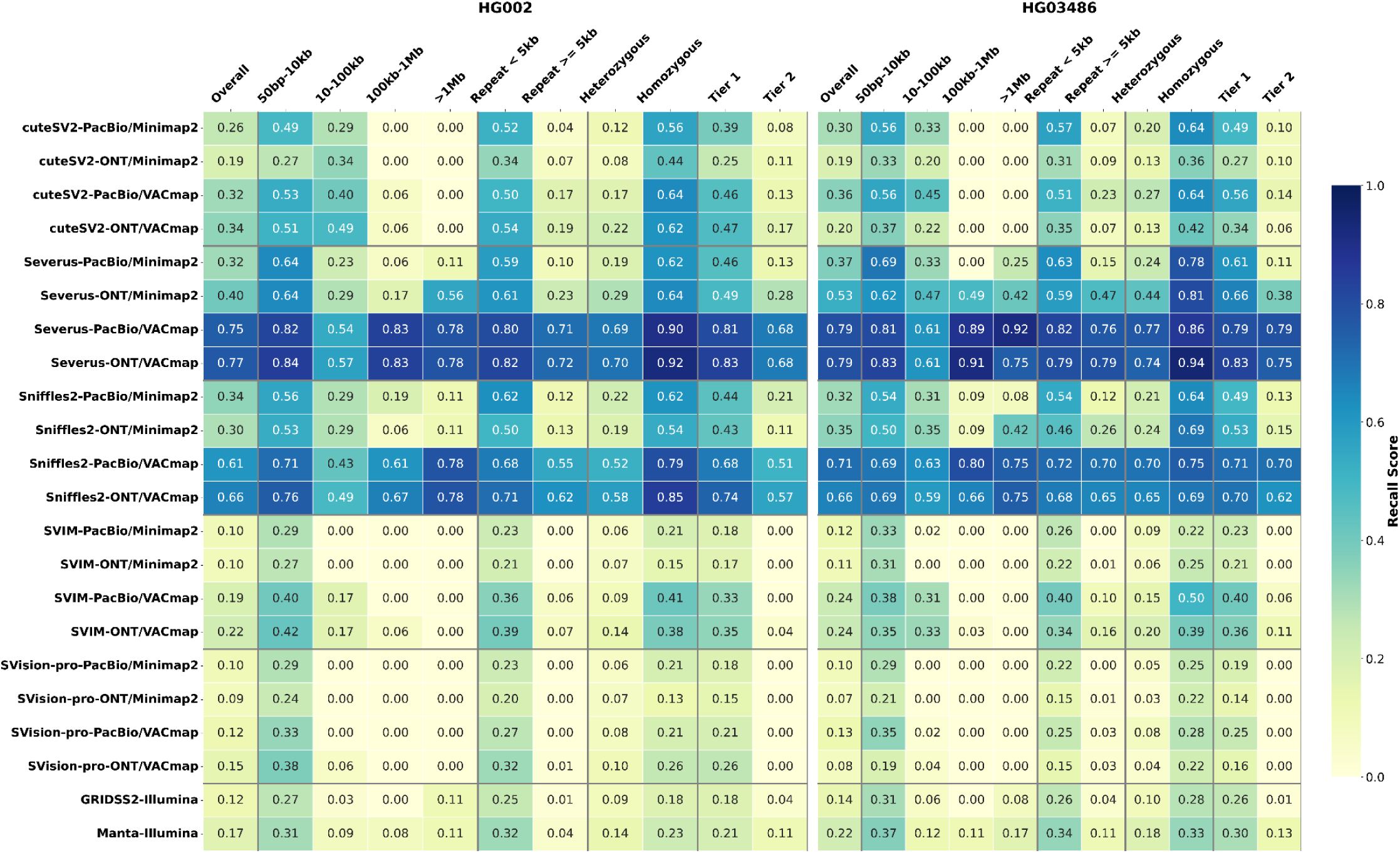
Comparative benchmarking of overall and stratified recall of inversion calling at 30× coverage for HG002 and HG03486 samples. Each row represents a specific combination of a structural variant (SV) caller, sequencing technology (PacBio or ONT), and alignment software (Minimap2 or VACmap). Short-read callers using Illumina data are included below. The columns display recall scores stratified by genomic context, including overall performance, variant size, repeat element length, zygosity (heterozygous vs. homozygous), and evidence-based confidence (Tier 1 vs. Tier 2). The color intensity of each cell corresponds to the recall score, with darker shades of blue indicating higher recall.

We first benchmarked the short-read callers on HG002. Overall, the recall was low, with two callers, GRIDSS2 and Manta, achieving 0.12 and 0.17, respectively. This is in contrast to long-read methods that were expected to perform better given the improved resolution in repeats. Upon closer investigation, we found the performance of long-read callers was highly aligner-dependent: with minimap2, top callers like Severus and Sniffles2 produced much higher sensitivity, e.g., in 30x ONT/minimap2, Severus reached a recall of 0.40, followed by Sniffles2 (0.30) and cuteSV2 (0.19), whereas SVIM and SVision-pro were markedly less sensitive (0.10 and 0.09). Replacing minimap2 with VACmap dramatically increased recall. Notably, the combination of Severus/Sniffles2 and VACmap consistently achieved the highest recall across a wide range of stratifications. For example, in 30x PacBio/VACmap data, Severus and Sniffles2 achieved the highest recalls (0.79 and 0.71), far above cuteSV2 (0.36), SVIM (0.24), and SVision-pro (0.13). cuteSV2 showed only limited recall gains with VACmap, consistent with a bias toward small events. For instance, in PacBio/VACmap data, the median length of true positive inversions called by cuteSV2, Severus, and Sniffles2 is 6.8 Kb, 239.9 Kb, and 209.3 Kb.

Stratified analyses revealed multiple consistent trends across all five samples assessed. First, Inversions flanked by short tandem repeats (<5 kb) were recovered with substantially higher recall than those flanked by longer repeats (≥5 kb). In 30x PacBio/Minimap2 data, Severus achieved a recall of 0.61 in short-repeat regions versus 0.23 in long-repeat regions. Second, VACmap further improved detection in challenging genomic contexts, including repeat-rich, heterozygous regions. For example, when switching from Minimap2 to VACmap, Sniffles2 recall in 30x ONT alignments improved from 0.13 to 0.62 in repeat-flanking loci and from 0.19 to 0.58 in heterozygous calls. Third, recall was also size-dependent, with the highest sensitivity for small inversions (50 bp-10 kb), followed by a gradual decline for larger variants, though sporadic increases were observed in the >1 Mb bin, which is likely due to limited representation in the benchmark set. Lastly, heterozygous inversions were consistently harder to detect than homozygous events (e.g., 30x Sniffles2 ONT/Minimap2: 0.19 vs. 0.54 recall). Thus, the recall overall shows already some interesting insights, but is of course limited, as we disregarded the precision and other factors so far. Nevertheless, this demonstrates further that the benchmark overall highlights several challenges for state-of-the-art SV callers and mapping methods.

### Performance evaluation across simple inversions

While the previous section focused on recall, precision is equally crucial for assessing variant caller performance. Therefore, we next evaluated the F1 score, which harmonizes both precision and recall into a single metric. To illustrate this analysis, we highlight results from HG002 and HG03486 evaluated on Tier 1 inversions in **Figure 4**, while comprehensive performance metrics across additional samples are provided in **Supplementary Figure 2**.

**Figure 4.**
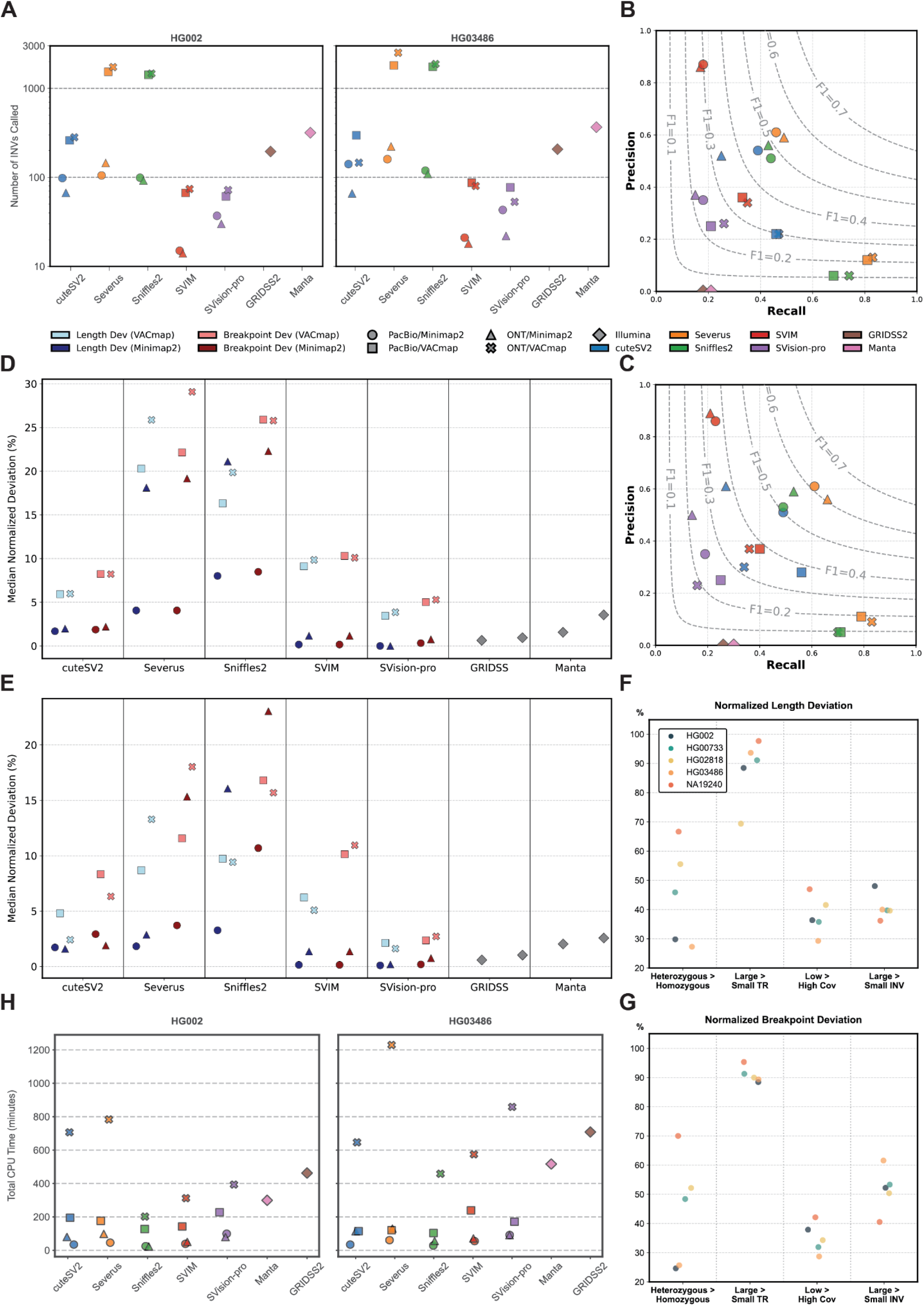
Comprehensive benchmarking of inversion-calling performance across SV callers on 30× HG002 and HG03486 long-read datasets on Tier 1 assembly-refined inversions. **(A)** Total number of inversions called by each tool under four sequencing/aligner workflows: PacBio/Minimap2 (circles), PacBio/VACmap (squares), ONT/Minimap2 (triangles), and ONT/VACmap (crosses). Illumina-based callers (GRIDSS2, Manta) are shown for comparison. **(B, C)** Precision-recall-f1 plots for inversion calls benchmarked against the Tier 1 truth sets for **(B)** HG002 and **(C)** HG03486. Each point represents the performance of a specific caller and technology/aligner combination, as detailed in the central legend. The concentric dashed lines are F1-score isoquants, indicating balanced precision-recall performance. **(D, E)** Median normalized deviation (%) in inversion length (blue colors) and breakpoint position (red colors) for each caller and each sequencing/aligner combination in HG002 **(D)** and HG03486 **(E)**. **(F, G)** Stratified analysis investigating the impact of genomic context on **(F)** normalized length deviation and **(G)** normalized breakpoint deviation. The barplots show the percentage of tests, across five human genomes (HG002, HG02818, HG03486, HG00733, and NA19240) with four hypotheses for increased deviation: 1) inversions flanked by large tandem repeats (Large > Small TR), 2) heterozygous inversions (Heterozygous > Homozygous), 3) large inversion size (Large > Small INV), and 4) low sequencing coverage (Low > High Coverage). A value of 100% on an axis indicates that the assumption is consistently held for that sample. **(H)** Total CPU time (minutes) required by each caller for HG002 (left) and HG03486 (right).

In the 30x HG002 sample, for the short-read callers, GRIDSS2 achieved an F1 score of 0.13, and Manta achieved an F1 score of 0.12 (**Figure 4B**). In contrast, long-read callers generally achieved a higher F1 score, resulting from higher precision and recall (**Figure 4A, B**). Again, we observed that the SV callers are highly dependent on the alignment method that is used. For example, in 30x ONT/minimap2, Severus reached the highest F1 score 0.53, followed by Sniffles2 (0.49), cuteSV2 (0.34), SVIM (0.28), and SVision-pro (0.22). Similar to recall, replacing minimap2 with VACmap dramatically increased the number of false positives and hence decreased the F1. For example, the F1 score dropped from 0.53 to 0.23 and from 0.49 to 0.12, for Severus and Sniffles2, respectively.

Besides, using VACmap data, SVIM improved to the highest F1 score with cuteSV2, Severus, SVision-pro, and Sniffles2 following behind. For example, in 30x HG002 PacBio/VACmap data, callers achieved F1 as 0.35 (SVIM), 0.30 (cuteSV2), 0.23 (SVision-pro), 0.21 (Severus), and 0.12 (Sniffles2). The steep F1 shifts exhibited by Sniffles2 and Severus reflect substantial recall gains offset by larger precision losses (**Supplementary Table 7**). Notably, SVIM and SVision-pro showed relatively stable F1 across callers, which relates to their low inversion calls despite the usage of aligners (**Figure 4B**).

Next, we investigated the number of inversions called by each caller with different aligners behind the F1 score (**Figure 4A**). It is interesting to note that short-read callers, such as GRIDSS2 and Manta, typically report a higher number of SV than long-read callers. For example, in the HG002 sample, GRIDSS2 and Manta called 195 and 317 inversions, respectively, while long-read callers called 14-145 inversions (e.g., Severus: 105-145, Sniffles2: 92-99). This is despite the higher recall on long-read callers. However, when we look at maximized recall for the long-read SV caller, using VACmap, we observe a dramatic increase in the reported inversions. For example, Severus and Sniffles2 exceeded 1,400 inversions per sample with VACmap, where Severus rose ∼15-fold in HG002 PacBio data (105 inversions with minimap2 to 1,533 with VACmap).

We then quantified normalized length and breakpoint deviations (**Methods; Figure 4D, E**) and integrated these with observed length distribution (**Supplementary Figure 3**). We found that long-read SV callers with minimap2 showed larger length and breakpoint deviations compared to short-read SV callers, whereas VACmap dramatically increased both deviations. These deviation patterns can be explained by the inversion length spectra and the increase in the number of true positive inversion calls. Short-read tools detected 17-23 true positives (TPs) and primarily detected events of 50 bp-10 kb, with Manta extending into the 10 kb-1 Mb range. Among long-read callers, minimap2-based calling detected 32-71 TPs and mostly concentrated in the 50 bp-100 kb range. In contrast, VACmap-based calling detected 54-173 TPs and allowed expansion into the ≥100 kb space, particularly for Severus and Sniffles2, which included hundreds of inversion calls exceeding 100 kb (**Supplementary Figure 3, Supplementary Table 7**). Among long-read SV callers (**Figure 4D, E**), SVision-pro had the smallest deviations in both aligner groups (e.g., SVision-pro: 0.00% length and 0.32% breakpoint deviation in 30x HG002 PacBio/minimap2), explained by its inversion calls being mostly below 10 Kb. SVIM was highly accurate with minimap2, yet had a large deviation with VACmap. For instance, with 30x HG002 PacBio data, SVIM’s normalized length and breakpoint deviations were just 0.16% with minimap2, but surged to 9.11% and 10.31% respectively, with VACmap. Sniffles2 and Severus showed the largest deviations with both aligners, indicating overall lower breakpoint accuracy, likely correlated with their higher true positive inversion numbers. For example, in 30x HG002 ONT/VACmap data, Sniffles2 and Severus generated deviations of 19.85%/25.86% (length), and 25.78%/29.07% (breakpoint), respectively.

This breakpoint accuracy investigation revealed a distinct hierarchy of effects (**Figure 4F, G**). Large flanking tandem repeats (>5 Kb) are the predominant factor contributing to larger deviations. In comparison, low coverage (10x), large inversion (>10 Kb), and heterozygosity demonstrated less influence (**Supplementary Table 3**).

Genotype concordance (gt-concordance) of Tier 1 inversions remained highly variable and sensitive to variant properties (**Supplementary Figure 4, 5**). In 30x datasets, Manta performed the best or the second-best in gt-concordance among all callers. Among long-read callers, SVIM achieved the highest overall gt-concordance, while cuteSV2 had the second-best genotyping ability. Sniffles2, Severus, and SVision-pro fell behind. GRIDSS2 was excluded as it didn’t perform genotyping of SVs. For instance, in 30x HG002 ONT/VACmap overall genotype concordance, SVIM and cuteSV2 achieved 0.88 and 0.78, respectively, while Manta had 0.86 with Illumina data. The further stratified analysis also indicated that large flanking TRs (>5 Kb), large inversion (>10 Kb), and heterozygosity will lead to lower genotype concordance. For example, in HG002 30x PacBio/minimap2 data, the genotype concordance of Severus and Sniffles2 dropped from 0.67 to 0.00 and from 0.39 to 0.25 when flanking TRs increased over 5 Kb. These results underscore that, even at high coverage, accurate genotyping of Tier 1 inversions remains a significant challenge across callers.

Sniffles2 was consistently the fastest caller, underscoring its suitability for population-scale studies (**Figure 1H**). In 30x HG002 PacBio/VACmap, Sniffles2 required 22.49 total CPU minutes, which was 3.5x and 4.4x faster than cuteSV2 (79.43 min) and Severus (98.06 min), respectively. Calling on minimap2 was uniformly faster than on VACmap, suggesting VACmap’s richer breakpoint landscape imposes computational overhead. Overall, long-read callers were faster than short-read tools (Supplementary Table 4).

Coverage subsampling (30x, 20x, 10x, 5x) confirmed that increased coverage generally improved F1 and recall (**Supplementary Figure 6; Supplementary Table 5**). In HG002, over 80% of results showed strong Pearson and/or Spearman correlations (≥0.7, p<0.05) for recall and F1. In Tier 1 inversions, aligner-specific patterns still hold. In minimap2 datasets, Severus and Sniffles2 maintained the highest F1 scores. For example, in HG002 ONT/minimap2 data, with descending 30x, 20x, 10x, and 5x coverage, Severus achieved 0.53, 0.55, 0.45, and 0.33, respectively, while Sniffles2 maintained stability across coverages at 0.49, 0.49, 0.47, and 0.34. In contrast, using VACmap data, SVIM and cuteSV2 stand out in F1 across high coverages. For instance, in HG002 PacBio/VACmap data, F1 scores of SVIM (0.35, 0.23, 0.11, 0.03) and cuteSV2 (0.30, 0.27, 0.13, 0.00) outperformed Severus (0.21, 0.21, 0.18, 0.13), with performance decreasing alongside coverage.

### Benchmarking of complex Inversions

We subsequently focused on the Tier 2 inversions as these represent complex structural variants (cSVs) which are composed of multiple breakpoints whose origin cannot be explained by a single end-joining or DNA exchange event[17,38,39]. These large, complex inversions can confound mapping and variant callers, leading to missed or imprecise calls. To account for this, we focused more on evaluating the presence of the inversion/cSV rather than the breakpoint resolution (**Methods**).

We first investigated the performance of Tier 2 inversions on 30x HG002 data. **Figure 5A** shows a ∼140 Kb Strand-seq indicated inversion (chr1:16653301-16795670) on chromosome 1 (1p36.13) flanked by large tandem repeats, where two separated assembly-refined heterozygous inversions were created when the complex breakpoints stopped the pipeline from merging two haplotypes into one homozygous representation. Only Severus and Sniffles2 detected inversions. However, both callers reported multiple, fragmented inversions for what appears to be a single complex event. This suggests a tendency for over-calling in regions with repetitive elements. The underlying complexity is visible in the ONT/VACmap alignment, which shows multiple nested breakpoints and inverted reads. While these calls were classified as true positives, their fragmented nature and deviated breakpoints highlight the difficulty in accurately defining SVs in such loci. **Figure 5B** shows a ∼39 Kb Strand-seq indicated inversion (chr18:20562340-20601393) located in the centromere of chromosome 18 (18q21.1), where the two haplotypes exhibit significant divergence in their assemblies, leading to two separated assembly-refined heterozygous inversions. In this highly complex region, Severus was the only tool to make a call. It reported three true positives and three false positives SVs, underscoring the reduced accuracy and high false-positive rate when calling cSVs within the centromere. The presence of multiple breakpoints and inverted reads in the alignment file further indicates a complex rearrangement that challenges current SV callers.

**Figure 5.**
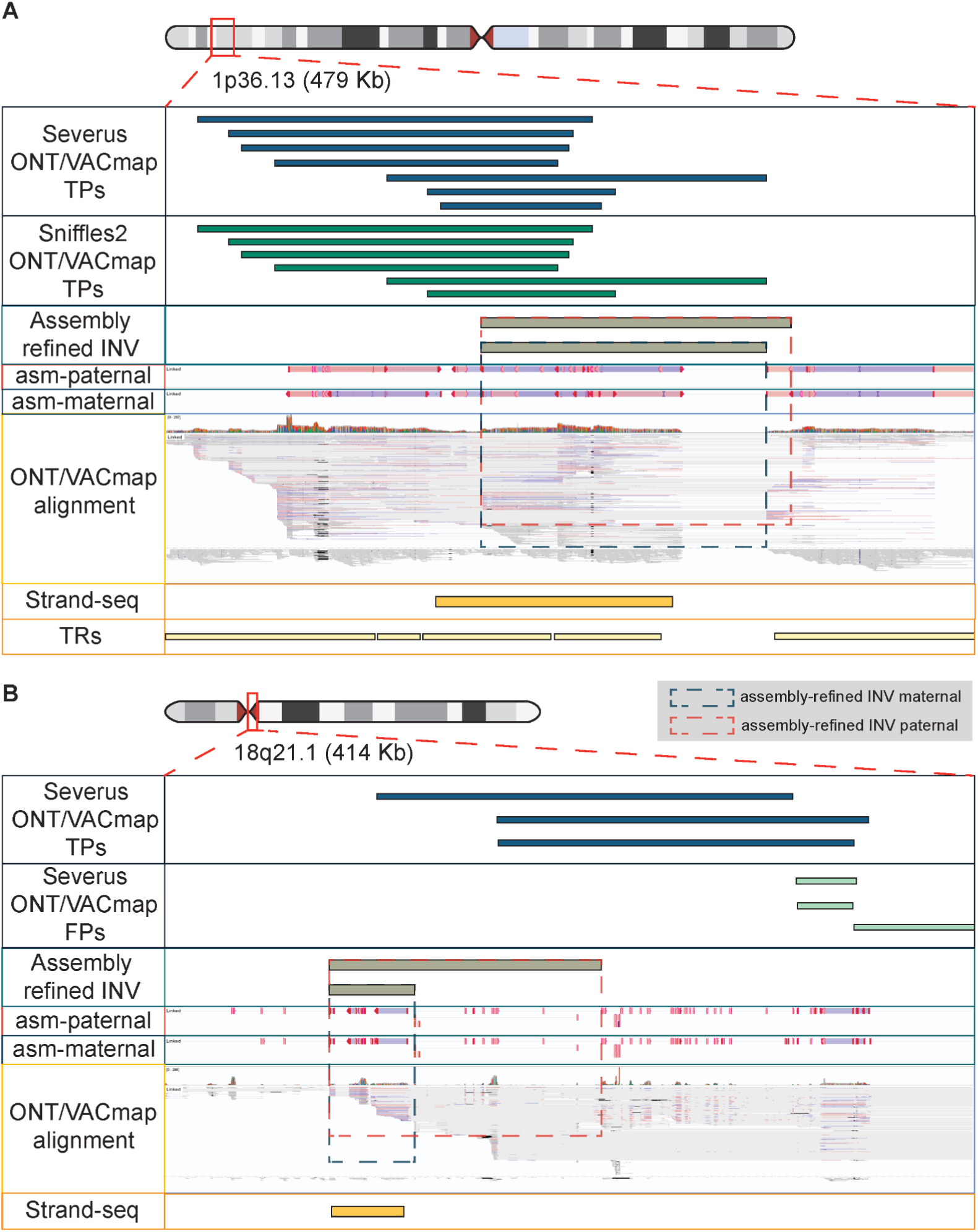
Performance of SV callers on inversions in complex genomic regions. **(A)** A ∼170 Kb inversion on chromosome 1 (1p36.13) flanked by large tandem repeats (TRs). Tracks from top to bottom show: True positive (TP) inversion calls from Severus and Sniffles2 on the ONT/VACmap data; the assembly-refined inversion; paternal and maternal haplotype assemblies (asm-paternal, asm-maternal); ONT/VACmap read alignments, where colored reads (blue and pink) are linked reads indicating inverted orientation, revealing multiple breakpoints; the Strand-seq inversion call confirming the event; and the location of the flanking tandem repeats. **(B)** A ∼39 Kb inversion on chromosome 18 (18q21.1) located within the centromere. Tracks from top to bottom show: True positive (TP) and false positive (FP) inversion calls from Severus; the two separated, assembly-refined heterozygous inversions from the divergent maternal and paternal haplotypes; paternal and maternal haplotype assemblies (asm-paternal, asm-maternal); ONT/VACmap read alignments showing complex signatures; and the corresponding Strand-seq inversion call.

Overall, the benchmark results showed that, although no SV caller fully captured all Tier 2 inversions, long-read callers outperform short-read callers on complex inversions (**Supplementary Fig. 7B, D**). For example, the best-performing long-read tools, Sniffles2 and Severus, achieved roughly 50-68% recall of known Tier 2 inversions at 30x with VACmap. In contrast, short-read callers, Manta or GRIDSS, recalled only a very small fraction (often under 10%). The universal low F1 scores revealed the complexity of the complex inversion calling: although Severus consistently achieved the highest F1 across samples, the numbers are still low. For example, for F1 score in 30x HG002 PacBio/VACmap data, Severus: 0.24, Sniffles2: 0.10, cuteSV2: 0.06, SVIM: 0.0, SVision-pro: 0.0, GRIDSS2: 0.03, Manta: 0.04.

There are several key observations in Tier 2 variants. First, Severus and Sniffles2 achieved the highest sensitivity even with low coverage data. cuteSV2 showed intermediate performance (lower recall than Sniffles2/Severus but higher than others), whereas SVIM and SVision-Pro performed poorly on complex inversions. Second, VACmap can lead to higher recall but lower precision and higher deviations compared to minimap2.

## Discussion

Inversions represent one of the most biologically meaningful yet systematically under-characterized classes of structural variation (SV). Although long-read sequencing and Strand-seq have accelerated our ability to detect these events[1,19], reliable benchmarking has lagged behind, focusing primarily on insertions and deletions[40–43]. The absence of a benchmark set has limited the development, optimization, and adoption of inversion callers, particularly for complex, repeat-flanked, or multi-breakpoint events[44,45]. Here, we address this gap by presenting a curated, multi-sample benchmark that integrates Strand-seq phasing, long-read assemblies, and rigorous breakpoint refinement. This resource establishes a consistent framework for evaluating inversion performance across technologies, aligners, and variant callers.

Our results reveal clear strengths and limitations of current approaches. For simple inversions in mappable regions, long-read methods have matured: callers such as Sniffles2 achieve high precision and recall, especially when combined with minimap2. However, performance drops substantially for variants embedded in segmental duplications or those spanning complex rearrangements, consistent with the known challenges of alignment ambiguity and breakpoint representation. We find that tools leveraging richer local signals (e.g., VACmap) improve recall in repetitive regions, but often at the cost of precision. Likewise, incorrect genotyping of heterozygous complex inversions remains common, emphasizing that detection alone is not sufficient and accurate allele assignment is still an open challenge. Most notably, no current method robustly resolves complex inversions, particularly nested or compound structures. These variants account for a non-trivial fraction of inversion diversity in human genomes and are increasingly implicated in disease and genome stability. Their persistent ambiguity underscores the need for algorithmic innovation. Both are important as we often face more complex SVs in disease backgrounds such as *MECP2[7,43,46]*. Clearly, there are also downsides to our proposed benchmark. First, there is a risk of assigning wrong genotypes to our benchmark inversions, especially in noisy regions, as not accurately accounting for two overlapping heterozygous inversions as one homozygous inversion remains a risk. Second, our breakpoint refinement, while performed across two independent aligners, cannot guarantee to provide the absolutely correct breakpoints, especially in Tier2 complex SV classes. One has to acknowledge that complex SV breakpoints are in highly repetitive regions (i.e., segmental duplications), which sometimes makes the breakpoint position arbitrary. We accounted for the evaluation of Tier2 inversion by only measuring presence and absence. However, it’s desirable to pinpoint these breakpoints accurately.

Besides the shortcomings of methods, there are multiple important key messages from this benchmark. People need to understand that SV callers rely heavily on correct alignments. While some default methods (e.g., minimap2 shown here but not exclusively) have reduced performance in highlighting inversions or complex rearrangements, others (e.g., VACmap) improve among them but currently may lead to many false positives genome-wide. This might be resolved with better adaptations of callers to novel alignment methods, co-developing aligners with SV callers[3], or forming hybrid alignment methods[47] that combine the best of both. For SV callers, outlining clear failure modes and genome contexts where performance declines provides actionable guidance for tool developers and users. Practically, our results suggest that researchers analyzing human genomes today should (i) trust simple inversions in non-repetitive sequence; (ii) treat inversion calls in segmental duplications with caution unless supported by orthogonal evidence; and (iii) avoid over-interpretation of complex inversion calls without local assembly or Strand-seq support. In all cases, we want to emphasize that visualisation of inversions or all types of SV often helps assess the fidelity of SV.

Looking forward, we anticipate continued convergence of sequencing modalities, graph methods, and machine-learning-driven refinement. Investments in benchmarking resources, including those focused on complex variation, will accelerate these developments and ultimately close the current gap between inversion discovery and biological interpretation. By releasing this resource to the community, we aim to provide a foundation for the next generation of inversion callers and to promote rigorous, reproducible structural variant analysis across large cohorts and clinical studies.

## Methods

### Data used in this study

The data used in this study are listed in **Supplementary Table 6** and the Data Availability section. The results of Strand-seq indicated inversions of 5 samples (HG002, HG02818, HG03486, HG00733, NA19240) come from a previous study[16]. For the HG002 sample, the raw long-read ONT data were downloaded from the ONT website (https://epi2me.nanoporetech.com/giab-2025.01/). We combined two runs (PAW70337 and PAW71238) using SAMtools (version 1.21) merge (see data availability). The HG002 PacBio data was downloaded from GIAB, while the short-read Illumina data came from NIST (SRA: SRR8861483). For the HG00733 sample, the raw long-read data, both PacBio HiFi and ONT Ultra-long, come from HGSVC, and the raw short-read Illumina data come from 1KG (SRA: ERP010495). For the NA19204 sample, the raw long-read PacBio HiFi data come from NCBI (SRA: SRR11363957), and ONT R10 data come from the HPRC consortia. The short-read Illumina data come from 1KG (SRA: ERP010495). For HG02818 and HG03486 samples, all the short-read Illumina data and long-read data, both PacBio HiFi and ONT R10, come from HPRC. All haplotyped-resolved assembly data come from the HPRC consortia.

### Assembly-refined Inversion Benchmark Set

A multi-step computational pipeline was designed to refine an initial set of Strand-seq inversion loci into a high-confidence, assembly-based callset. The entire workflow was performed independently on haplotype-resolved assembly BAMs generated by Minimap2 and VACmap before the final results were merged.

Haplotype-Specific Breakpoint Detection: For each input Strand-seq indicated locus, precise breakpoints were identified from paternal and maternal BAM files through a custom detection process. This process searched an expanded window (300% of inversion length, max 3 Mb) around each locus for reads indicating structural variation. Candidate breakpoints were extracted from reads with a mapping quality (MAPQ) ≥ 10 that either contained a supplementary alignment (SA tag) or exhibited soft-clipping of ≥10 bp. These raw positions were clustered if within 100 bp of each other, and the optimal pair of start and end breakpoint clusters was selected based on a weighted composite score evaluating proximity to the initial BED locus (60%), opposite-strand signal clarity (10%), read support (15%), and length similarity (15%). Each call was assigned a FILTER status based on its precision, defined by the ratio of the breakpoint cluster’s genomic range to the final inversion length (SVLEN). A call was marked PASS_PRECISE if this ratio was ≤ 0.20; PASS_IMPRECISE if the ratio was > 0.20 and ≤ 0.50, or if high ambiguity was detected (≥4 unique clusters per side or ≥5 valid cluster pairs); and LOWCONF if the ratio was > 0.50.

Diploid Genotype Merging: The resulting haplotype-specific calls were grouped by their ORIGINAL_LOCUS identifier and merged into diploid genotypes. If both haplotypes yielded a signal and their start/end breakpoints were concordant (differing by ≤500 bp), they were merged into a single record with a homozygous genotype (1/1). These merged calls were assigned a MERGED_HAP INFO tag to stratify confidence: PRECISE_CONF for identical precise calls or one precise call against a NOSIGNAL haplotype; PRECISE_Mid_CONF for precise calls differing by ≤500 bp; and IMPRECISE_Two_Hap for two concordant imprecise calls. If precise calls differed by >500 bp, they were labeled PRECISE_Low_CONF and kept as separate heterozygous (0/1) records. Variants with only one haplotype signal were also kept as heterozygous calls and labeled IMPRECISE_One_NoSignal if the call was imprecise.

Redundant Inversion Filtering and Repeat Annotation: To remove nested artifacts, the callset was filtered by first parsing the initial BED file to identify any inversion strictly contained within the coordinates of a larger inversion on the same chromosome. The identifiers of these contained loci were used to create a regex pattern to filter them from the VCF file. All loci on chrY were also removed during this step. The filtered VCF was then annotated by querying reference BED files for tandem repeats from the adotto database[48] and centromeres annotation from UCSC[49,50] using an interval-tree-based approach. Variants were flagged in the INFO field as Repeat_ge5Kb or Repeat_ge10Kb if their breakpoints overlapped with segmental duplications of corresponding size, and a Centromere flag was added if the inversion interval overlapped with known centromeric regions.

Hierarchical Merging of Aligner Callsets: The final curated callset was created by merging the two processed VCFs (one from Minimap2, one from VACmap) through a hierarchical process. First, it excluded any locus previously annotated with the Centromere flag or located on chrY. For the remaining loci, the Minimap2-derived call was selected if it was not NOSIGNAL, as two aligners performed similarly in relatively simple regions. If the Minimap2 call was NOSIGNAL, the corresponding VACmap-derived call was used as a fallback, as VACmap’s ability to detect complex rearrangements. Loci where both aligners reported NOSIGNAL were discarded.

Callset Characterization and Comparison: The final VCF was analyzed to generate summary statistics on variant categories, size distributions, and overlap with genomic features. A detailed comparison between the two aligner-specific callsets (pre-merge) was conducted by generating a confusion matrix that compares call categories for each shared ORIGINAL_LOCUS to assess concordance and complementarity.

### Overview of benchmarked methods

#### Mapping methodology

BWA (v0.7.17) was run on paired-end Illumina short-read data with -M and -R parameters to mark shorter split hits as secondary and add read group header line, respectively. Long-read sequencing data were aligned by minimap2 and VACmap. Minimap2 (v2-2.28) was run with sequencing technology-specific parameters (PacBio: -Y -ax map-hifi -H, ONT: -Y -ax map-ont) with -R and -MD options to add read group line and the MD tag, respectively. While VACmap (v1.0.1) was run in mode S to increase sensitivity in variant detection, --MD, --rg-id, --rg-sm parameters were used to add MD tag and add read group line. Resulting alignments were converted to BAM format, sorted, and indexed using SAMtools (version 1.21).

#### Inversion calling methodology

All callers were run under the default or recommended parameters. For short-read data, Manta (v1.6.0) was used to call SVs with default settings following a Manta embedded script, convertInversion.py, to convert breakpoints into inversions. GRIDSS (v2.13.2) was used to call BNDs with default settings and a recommended exclude list for the GRCh38 reference genome. An embedded script was then performed to convert BNDs into inversion-only VCF files. For long-read callers, most were run in default parameters as well. cuteSV2 (v2.1.1) was used to call inversions based on the recommended parameter settings for PacBio HiFi and ONT data. For PacBio HiHi data, max_cluster_bias_INS was set to 1000, diff_ratio_merging_INS was set to 0.9, max_cluster_bias_DEL was set to 1000, and diff_ratio_merging_DEL was set to 0.5. In the ONT data, those parameters were set to 100, 0.3, 100, 0.3, respectively. Severus (v1.1) was run in germline mode with default parameters and the recommended VNTR annotation file for the GRCh38 reference genome. Due to a known issue in version 1.1, where the END tag was missing from the INFO field of inversion records in the output VCF, an in-house script was used to append the missing END annotations to enable downstream breakpoint analyses. Sniffles2 (v2.7.0) was run in germline mode with default settings. SVIM (v2.0.0) does not include filtering steps in its main pipeline, and we were not able to identify a recommended default cutoff for the quality value that SVIM outputs along with its SV calls. Therefore, in line with previous SV caller benchmarks, we filtered the output of SVIM to include only calls with a minimum read support of 10 by default (equal to the default of cuteSV2 and Sniffles2). SVision-pro (v1.8) was run in germline mode using the model_liteunet_256_8_16_32_32_32 model and the BED file recommended by the authors to exclude centromeric and heterochromatic regions. Because SVision-pro outputs complex SV annotations using concatenated labels (e.g., SVTYPE=INS+dDUP+DEL), an in-house script was used to extract any SV records that include inversion components for benchmarking.

### Truvari bench parameters

In our analysis, we executed truvari bench --pctseq 0 --pick multi --chunksize ${size_dependent} --refdist ${size_dependent} --pctsize 0.3 -b $ref_vcf -c $input_vcf --sizemax 5400000 to rigorously evaluate called INVs by each caller against the assembly-refined truth set in Truvari v5.2.0. For small variants ranging from 50 bp to 10 kb, we required high positional accuracy by setting the maximum allowed reference distance (--refdist) and the genomic comparison window (--chunksize) to 1 kb. For medium-sized variants from 10 kb to 100 kb, these parameters were increased to 10 kb. To accommodate the greater ambiguity in the breakpoint coordinates of large-scale events, a more lenient window of 100 kb was used for all variants larger than 100 kb.

Across all size-stratified analyses, a consistent framework of parameters was applied to ensure a uniform and fair comparison. We focused the evaluation on positional and size accuracy by disabling the sequence similarity requirement (--pctseq 0) while setting a lenient reciprocal size similarity threshold (--pctsize 0.3) requiring 30% reciprocal overlap in variant lengths, accommodating comparisons of small to large events without excluding low-similarity calls. We also constrained the analysis to variants no larger than 5.4 Mb with --sizemax 5400000 as the largest inversion from Strand-seq is ∼5.3 Mb, filtering out exceedingly large or potentially artifactual calls.

Absolute deviations in inversion length and breakpoints were computed from Truvari’s output fields. Absolute length deviation was the absolute value of SizeDiff, and absolute breakpoint deviation was the sum of the absolute values of StartDistance and EndDistance. To account for inversion size, normalized deviations were obtained by dividing each absolute deviation by the called inversion length (caller_inv_len) where normalized length deviation = abs(SizeDiff)/caller_inv_len, and normalized breakpoint deviation = [abs(StartDistance) + abs(EndDistance)]/caller_inv_len. The sample-wise deviations were calculated as the median of normalized values of all true positive inversions. The running time for each tool was recorded as the total CPU time consumed during execution.

### Computer specifications

All tests were performed in a high-performance cluster with Intel Xeon Gold 6148 CPU @ 2.40 GHz; the memory allocation was 32 GB unless otherwise stated; and the number of CPU cores allocated was eight unless otherwise stated. All CPU time is given as the sum of all compute times as if a single core were used. We measured and reported the total CPU time and wall clock time using the UNIX time command.

## Data availability

The data used in this study are listed in the **Supplementary Table S6.**

All VCF files representing assembly-based benchmark sets used in this study were uploaded to Zenodo: https://zenodo.org/records/17624133

All VCF files of SV callers across datasets and coverages can be found at GitHub *data_zenodo* section: https://github.com/jamesc99/INV-Benchmark/tree/main/data_zenodo

## Code availability

Scripts for the analysis are available at https://github.com/jamesc99/INV-Benchmark/

## Acknowledgments

FJS and SC are partially supported by NIH grants (1UG3NS132105 and 1U01HG011758-01)

## Contributions

S.C. performed study design, pipeline development, data analysis, and contributed to writing. F.J.S. conceived and supervised the study, led project planning, and contributed to writing the manuscript. All authors reviewed and approved the final manuscript.

## References

1. Mahmoud M, Gobet N, Cruz-Dávalos DI, Mounier N, Dessimoz C, Sedlazeck FJ. Structural variant calling: the long and the short of it. Genome Biol. 2019;20:246.

2. Smolka M, Paulin LF, Grochowski CM, Horner DW, Mahmoud M, Behera S, et al. Detection of mosaic and population-level structural variants with Sniffles2. Nat Biotechnol. 2024;42:1571–80.

3. Sedlazeck FJ, Rescheneder P, Smolka M, Fang H, Nattestad M, von Haeseler A, et al. Accurate detection of complex structural variations using single-molecule sequencing. Nat Methods. 2018;15:461–8.

4. Carvalho CMB, Lupski JR. Mechanisms underlying structural variant formation in genomic disorders. Nat Rev Genet. 2016;17:224–38.

5. Bansal V, Bashir A, Bafna V. Evidence for large inversion polymorphisms in the human genome from HapMap data. Genome Res. 2007;17:219–30.

6. Bondeson M-L, Dahl N, Malmgren H, Kleijer WJ, Tönnesen T, Carlberg B-M, et al. Inversion of the IDS gene resulting from recombination with IDS-related sequences in a common cause of the Hunter syndrome. Hum Mol Genet. 1995;4:615–21.

7. Pehlivan D, Huang C, Harris HK, Coquery C, Mahat A, Maletic-Savatic M, et al. Comprehensive assessment reveals numerous clinical and neurophysiological differences between MECP2-allelic disorders. Ann Clin Transl Neurol. 2025;12:433–47.

8. David D, Morais S, Ventura C, Campos M. Female haemophiliac homozygous for the factor VIII intron 22 inversion mutation, with transcriptional inactivation of one of the factor VIII alleles. Haemophilia. 2003;9:125–30.

9. Tizzano EF, Cornet M, Domènech M, Baiget M. Exclusion of mosaicism in Spanish haemophilia A families with inversion of intron 22. Haemophilia. 2003;9:584–7.

10. Kanto S, Chiba N, Tanaka Y, Fujita S, Endo M, Kamada N, et al. The PEBP2beta/CBF beta-SMMHC chimeric protein is localized both in the cell membrane and nuclear subfractions of leukemic cells carrying chromosomal inversion 16. Leukemia. 2000;14:1253–9.

11. Barresi GM, Albitar M, O’Brien SM. Acute myeloid leukemia, inversion 16, occurring in a patient with chronic lymphocytic leukemia. Leuk Lymphoma. 2000;38:621–5.

12. Fonseca H, da Silva TM, Saraiva M, Santolalla ML, Sant’Anna HP, Araujo NM, et al. Genomic regions 10q22.2, 17q21.31, and 2p23.1 can contribute to a lower lung function in African descent populations. Genes (Basel). 2020;11:1047.

13. Bosch N, Morell M, Ponsa I, Mercader JM, Armengol L, Estivill X. Nucleotide, cytogenetic and expression impact of the human chromosome 8p23.1 inversion polymorphism. PLoS One. 2009;4:e8269.

14. Pettersson M, Grochowski CM, Wincent J, Eisfeldt J, Breman AM, Cheung SW, et al. Cytogenetically visible inversions are formed by multiple molecular mechanisms. Hum Mutat. 2020;41:1979–98.

15. Balachandran P, Walawalkar IA, Flores JI, Dayton JN, Audano PA, Beck CR. Transposable element-mediated rearrangements are prevalent in human genomes. Nat Commun. 2022;13:7115.

16. Porubsky D, Höps W, Ashraf H, Hsieh P, Rodriguez-Martin B, Yilmaz F, et al. Recurrent inversion polymorphisms in humans associate with genetic instability and genomic disorders. Cell. 2022;185:1986–2005.e26.

17. Bilgrav Saether K, Eisfeldt J, Bengtsson J, Lun MY, Grochowski CM, Mahmoud M, et al. Mind the gap: the relevance of the genome reference to resolve rare and pathogenic inversions. medRxiv. 2024;2024.04.22.24305780.

18. Shen J, Cheng S, Purushotham D, Zhuo X, Du AY, Zhang W, et al. Exploring the epigenome profiles of repetitive elements with the WashU Repeat Browser. Genome Res [Internet]. 2025 [cited 2025 Feb 24]; Available from: https://pubmed.ncbi.nlm.nih.gov/39984199/

19. Sanders AD, Hills M, Porubský D, Guryev V, Falconer E, Lansdorp PM. Characterizing polymorphic inversions in human genomes by single-cell sequencing. Genome Res. 2016;26:1575–87.

20. Falconer E, Hills M, Naumann U, Poon SSS, Chavez EA, Sanders AD, et al. DNA template strand sequencing of single-cells maps genomic rearrangements at high resolution. Nat Methods. 2012;9:1107–12.

21. Ding H, Sedlazeck FJ, Liao Z, Pu L, Zhu S. VACmap: An accurate long-read aligner for unraveling complex genomic rearrangements [Internet]. bioRxiv. 2023. Available from: 10.1101/2023.08.03.551566

22. Zook JM, Catoe D, McDaniel J, Vang L, Spies N, Sidow A, et al. Extensive sequencing of seven human genomes to characterize benchmark reference materials. Sci Data. 2016;3:160025.

23. Zook JM, McDaniel J, Parikh H, Heaton H, Irvine SA, Trigg L, et al. Reproducible integration of multiple sequencing datasets to form high-confidence SNP, indel, and reference calls for five human genome reference materials [Internet]. bioRxiv. bioRxiv; 2018. Available from: 10.1101/281006

24. 1000 Genomes Project Consortium, Auton A, Brooks LD, Durbin RM, Garrison EP, Kang HM, et al. A global reference for human genetic variation. Nature. 2015;526:68–74.

25. Fairley S, Lowy-Gallego E, Perry E, Flicek P. The International Genome Sample Resource (IGSR) collection of open human genomic variation resources. Nucleic Acids Res. 2020;48:D941–7.

26. Liao W-W, Asri M, Ebler J, Doerr D, Haukness M, Hickey G, et al. A draft human pangenome reference. Nature. 2023;617:312–24.

27. Porubsky D, Harvey WT, Rozanski AN, Ebler J, Höps W, Ashraf H, et al. Inversion polymorphism in a complete human genome assembly. Genome Biol. 2023;24:100.

28. Li H. Minimap2: pairwise alignment for nucleotide sequences. Bioinformatics. 2018;34:3094–100.

29. Thorvaldsdóttir H, Robinson JT, Mesirov JP. Integrative Genomics Viewer (IGV): high-performance genomics data visualization and exploration. Brief Bioinform. 2013;14:178–92.

30. Cameron DL, Baber J, Shale C, Valle-Inclan JE, Besselink N, van Hoeck A, et al. GRIDSS2: comprehensive characterisation of somatic structural variation using single breakend variants and structural variant phasing. Genome Biol. 2021;22:202.

31. Chen X, Schulz-Trieglaff O, Shaw R, Barnes B, Schlesinger F, Källberg M, et al. Manta: rapid detection of structural variants and indels for germline and cancer sequencing applications. Bioinformatics. 2016;32:1220–2.

32. Jiang T, Liu S, Cao S, Wang Y. Structural variant detection from long-read sequencing data with cuteSV. Methods Mol Biol. 2022;2493:137–51.

33. Keskus AG, Bryant A, Ahmad T, Yoo B, Aganezov S, Goretsky A, et al. Severus detects somatic structural variation and complex rearrangements in cancer genomes using long-read sequencing. Nat Biotechnol. 2025;1–11.

34. Heller D, Vingron M. SVIM: structural variant identification using mapped long reads. Bioinformatics. 2019;35:2907–15.

35. Wang S, Lin J, Jia P, Xu T, Li X, Liu Y, et al. De novo and somatic structural variant discovery with SVision-pro. Nat Biotechnol. 2024;1–5.

36. Li H. New strategies to improve minimap2 alignment accuracy. Bioinformatics. 2021;37:4572–4.

37. English AC, Menon VK, Gibbs R, Metcalf GA, Sedlazeck FJ. Truvari: Refined structural variant comparison preserves Allelic diversity [Internet]. bioRxiv. 2022. Available from: 10.1101/2022.02.21.481353

38. Quinlan AR, Hall IM. Characterizing complex structural variation in germline and somatic genomes. Trends Genet. 2012;28:43–53.

39. Schuy J, Grochowski CM, Carvalho CMB, Lindstrand A. Complex genomic rearrangements: an underestimated cause of rare diseases. Trends Genet. 2022;38:1134–46.

40. Wagner J, Olson ND, Harris L, McDaniel J, Cheng H, Fungtammasan A, et al. Curated variation benchmarks for challenging medically relevant autosomal genes. Nat Biotechnol. 2022;40:672–80.

41. Zook JM, Hansen NF, Olson ND, Chapman L, Mullikin JC, Xiao C, et al. A robust benchmark for detection of germline large deletions and insertions. Nat Biotechnol. 2020;38:1347–55.

42. Daniels CA, Abdulkadir A, Cleveland MH, McDaniel JH, Jáspez D, Rubio-Rodríguez LA, et al. A robust benchmark for detecting low-frequency variants in the HG002 Genome In A Bottle NIST reference material [Internet]. bioRxivorg. 2024. Available from: 10.1101/2024.12.02.625685

43. Cheng S, Zhang Q, Zheng X, Jhangiani SN, Weir JC, Farek JR, et al. Constellation illuminates rare disease genetics [Internet]. medRxiv. 2025 [cited 2025 Oct 20]. p. 2025.10.15.25337675. Available from: https://www.medrxiv.org/content/10.1101/2025.10.15.25337675v1.abstract

44. Majidian S, Agustinho DP, Chin C-S, Sedlazeck FJ, Mahmoud M. Genomic variant benchmark: if you cannot measure it, you cannot improve it. Genome Biol. 2023;24:221.

45. Nardone GG, Andrioletti V, Santin A, Morgan A, Spedicati B, Concas MP, et al. A hitchhiker guide to structural variant calling: A comprehensive benchmark through different sequencing technologies. Biomedicines. 2025;13:1949.

46. Grochowski CM, Bengtsson JD, Du H, Gandhi M, Lun MY, Mehaffey MG, et al. Inverted triplications formed by iterative template switches generate structural variant diversity at genomic disorder loci. Cell Genom. 2024;4:100590.

47. Fu Y, Mahmoud M, Muraliraman VV, Sedlazeck FJ, Treangen TJ. Vulcan: Improved long-read mapping and structural variant calling via dual-mode alignment. Gigascience [Internet]. 2021;10. Available from: 10.1093/gigascience/giab063

48. English AC, Dolzhenko E, Ziaei Jam H, McKenzie SK, Olson ND, De Coster W, et al. Analysis and benchmarking of small and large genomic variants across tandem repeats. Nat Biotechnol [Internet]. 2024; Available from: 10.1038/s41587-024-02225-z

49. Bailey JA, Gu Z, Clark RA, Reinert K, Samonte RV, Schwartz S, et al. Recent segmental duplications in the human genome. Science. 2002;297:1003–7.

50. Bailey JA, Yavor AM, Massa HF, Trask BJ, Eichler EE. Segmental duplications: organization and impact within the current human genome project assembly. Genome Res. 2001;11:1005–17.

