## Supplementary Figures for "Benchmark for simple and complex genome inversions"

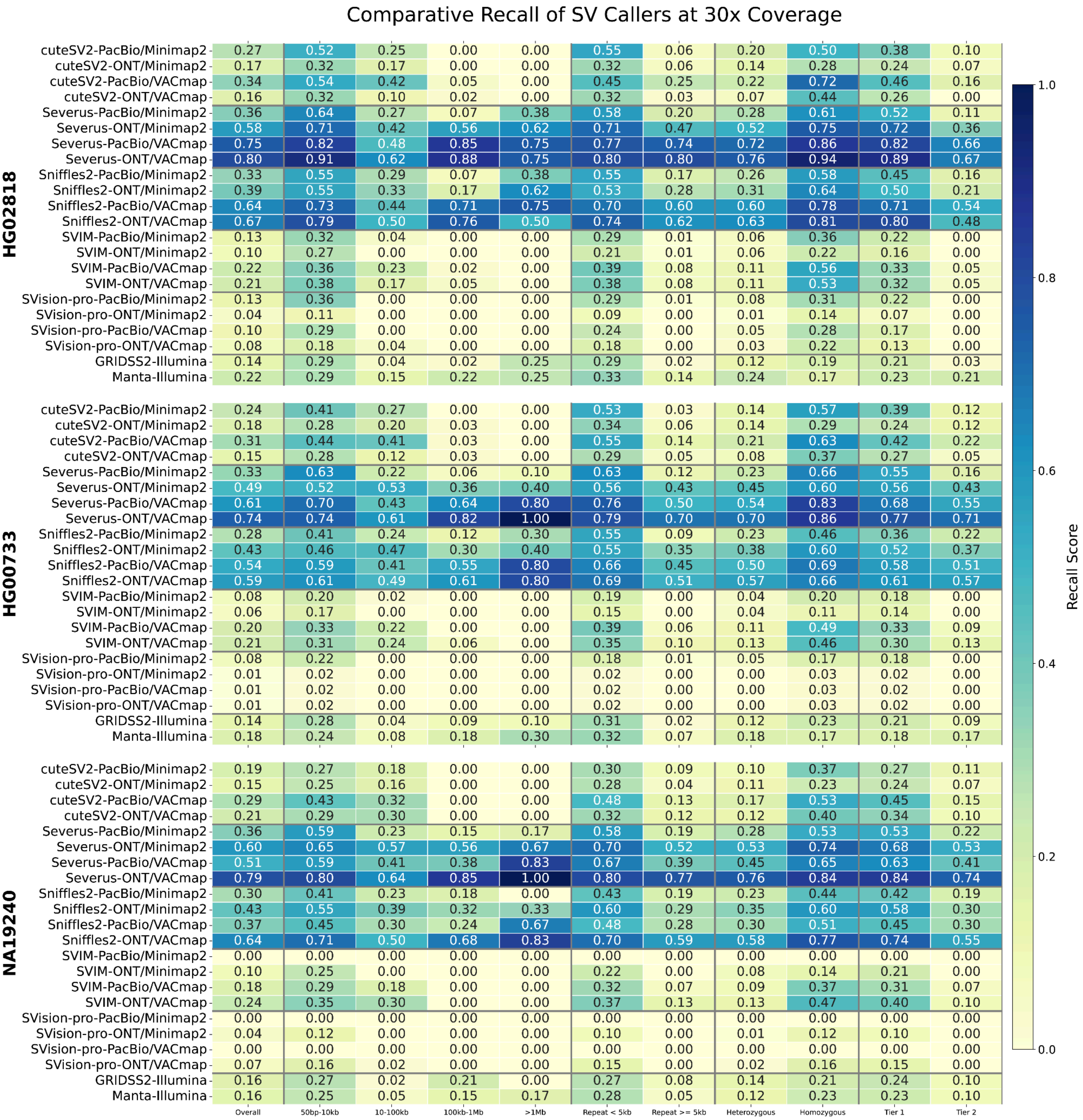


***Supplementary Figure 1: Comparative benchmarking of overall and stratified recall of inversion calling at 30× coverage for HG00733, HG02818, and NA19240 samples.*** *Each row represents a specific combination of a structural variant (SV) caller, sequencing technology (PacBio or ONT), and alignment software (Minimap2 or VACmap). Short-read callers using Illumina data are included below. The columns display recall scores stratified by genomic context, including overall performance, variant size, repeat element length, zygosity (heterozygous vs. homozygous), and evidence-based confidence (Tier 1 vs. Tier 2). The color intensity of each cell corresponds to the recall score, with darker shades of blue indicating higher recall.*

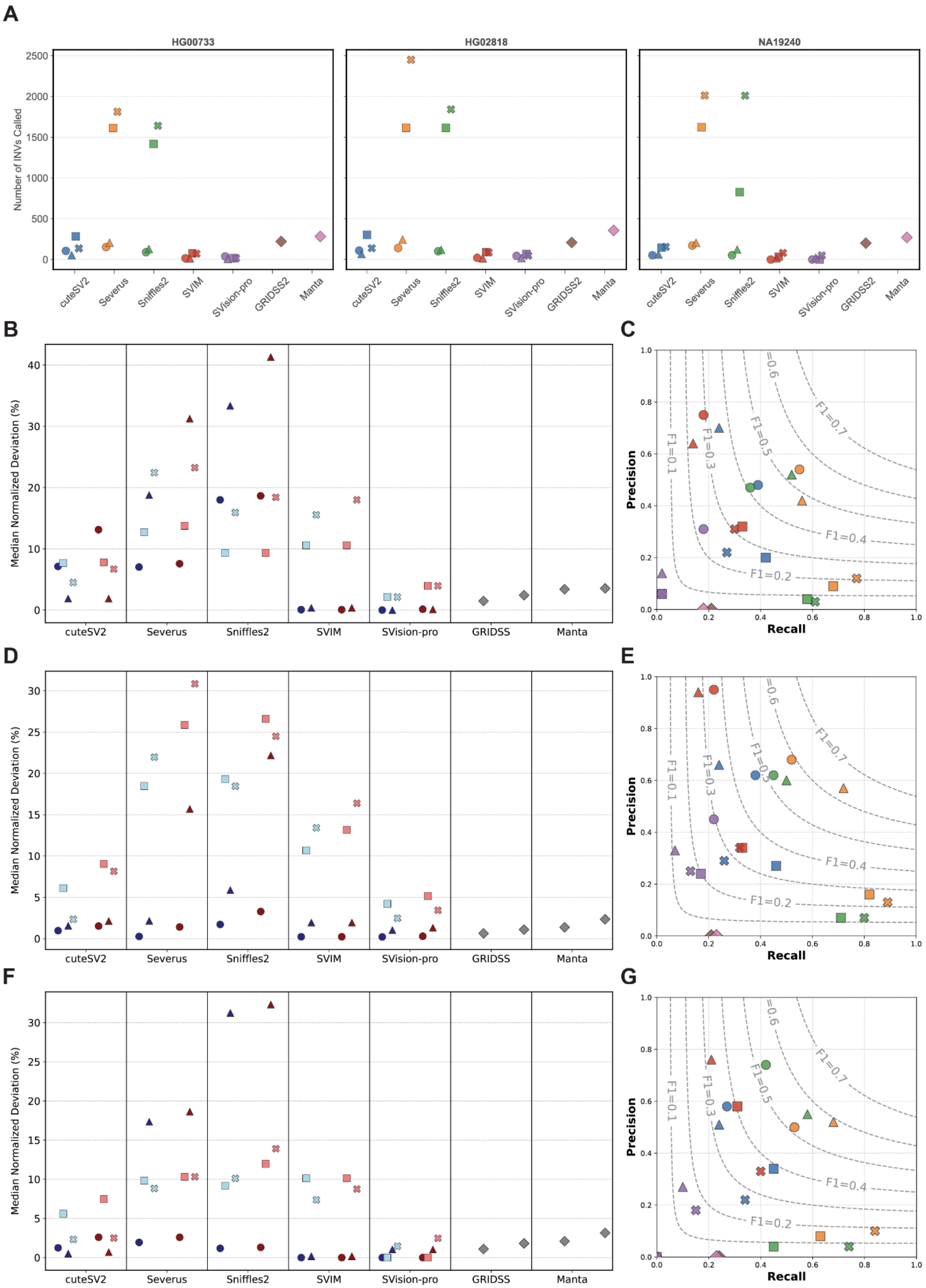

A
HG00733
HG02818
NA19240
Number of INVs Called
cuteSV2
Severus
Sniffles2
SVIM
SVision-pro
GRIDSS2
Manta
B
C
D
E
F
G
Median Normalized Deviation (%)
GRIDSS
Precision
Recall
F1=0.1
F1=0.2
F1=0.3
F1=0.4
F1=0.5
F1=0.6
F1=0.7

***Supplementary Figure 2: Tier 1 inversion calling results on 30x HG00733, HG02818, and NA19240 samples. (A)*** *Total number of inversions called by each tool under four sequencing/aligner workflows: PacBio/Minimap2 (circles), PacBio/VACmap (squares), ONT/Minimap2 (triangles), and ONT/VACmap (crosses). Illumina-based callers (GRIDSS2, Manta) are shown for comparison.* ***(B, D, F)*** *Median normalized deviation (%) in inversion length (blue colors) and breakpoint position (red colors) for each caller and each sequencing/aligner combination in HG00733* ***(B),*** *HG02818* ***(D)****, and HG03486* ***(F). (C, E, G)*** *Precision-recall-f1 plots for inversion calls benchmarked against the Tier 1 truth sets for HG00733* ***(C),*** *HG02818* ***(E)****, and HG03486* ***(G)****. Each point represents the performance of a specific caller and technology/aligner combination, as detailed in the central legend. The concentric dashed lines are F1-score isoquants, indicating balanced precision-recall performance.*

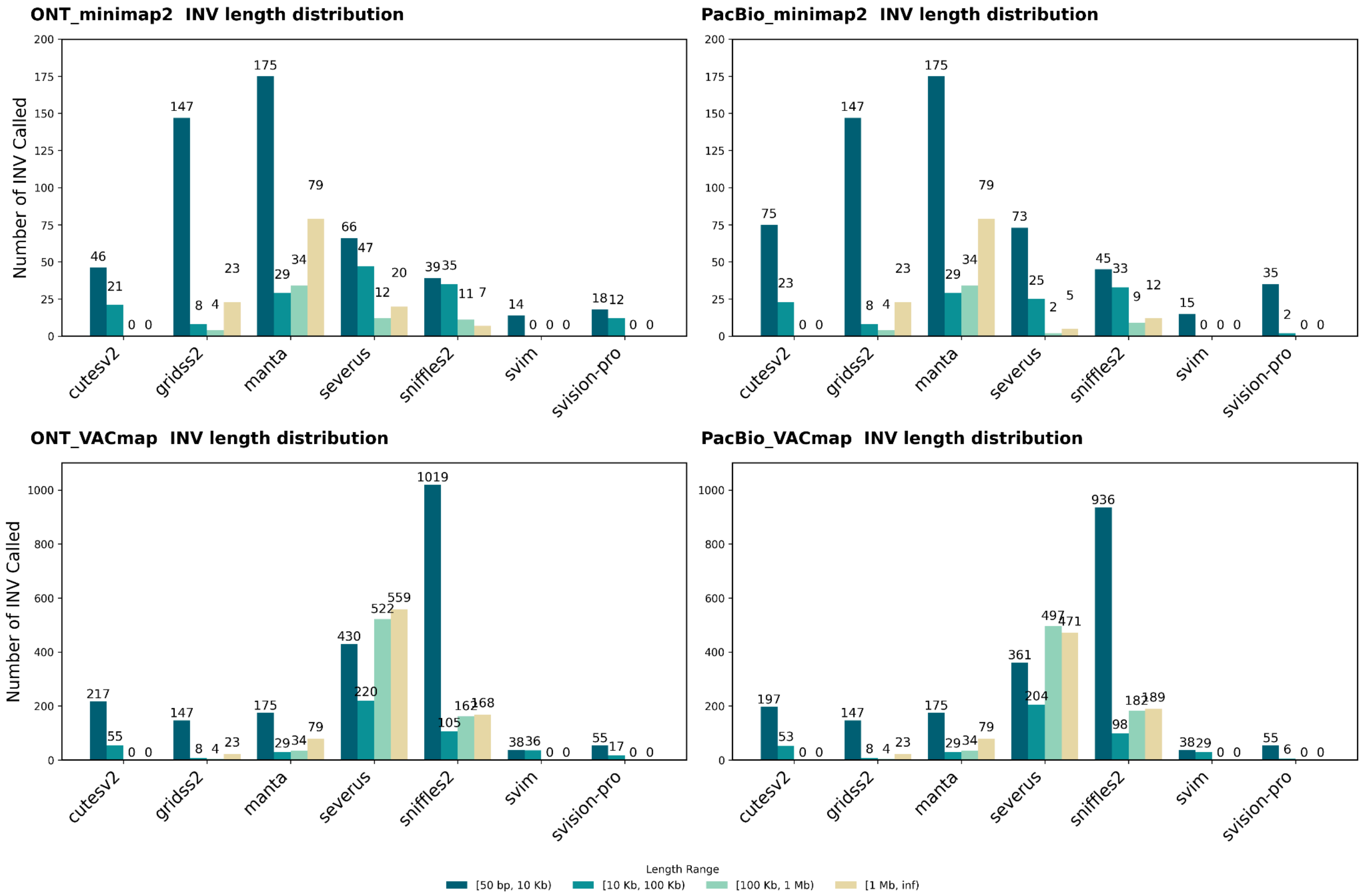


***Supplementary Figure 3: Length distribution of inversions on 30x HG002 data on the GRCh38 reference genome.*** *The data is stratified into four panels based on the sequencing platform and aligner used.* ***Top Left:*** *PacBio (PB) data aligned with Minimap2.* ***Top Right:*** *PacBio (PB) data aligned with VACmap.* ***Bottom Left:*** *Oxford Nanopore (ONT) data aligned with Minimap2.* ***Bottom Right:*** *Oxford Nanopore (ONT) data aligned with VACmap. In each panel, the X-axis represents five callers (Severus, Sniffles2, cuteSV2, SVision-pro, and SVIM), while the Y-axis indicates the count of variants. Inversion lengths are stratified into four categories: 50bp-10Kb, 10Kb-100Kb, 100Kb-1Mb, and larger than 1Mb.*

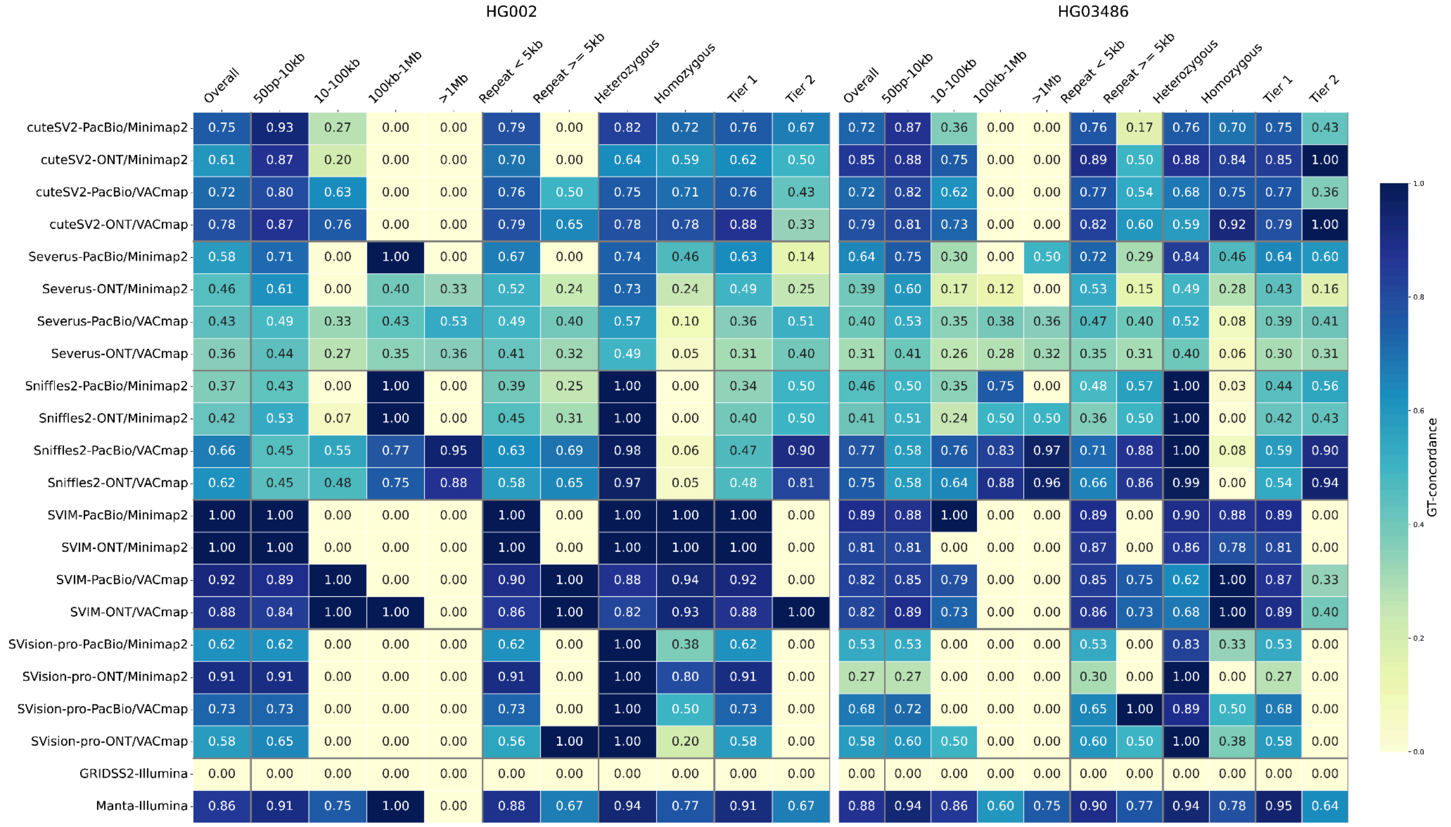


***Supplementary Figure 4: Genotype concordance of SV callers at 30x coverage of HG002 and HG03486 samples.*** *Each row represents a specific combination of a structural variant (SV) caller, sequencing technology (PacBio or ONT), and alignment software (Minimap2 or VACmap). Short-read callers using Illumina data are included below. The columns display GT-concordance scores stratified by genomic context, including overall performance, variant size, repeat element length, zygosity (heterozygous vs. homozygous), and evidence-based confidence (Tier 1 vs. Tier 2). The color intensity of each cell corresponds to the GT-concordance, with darker shades of blue indicating higher genotyping performance.*

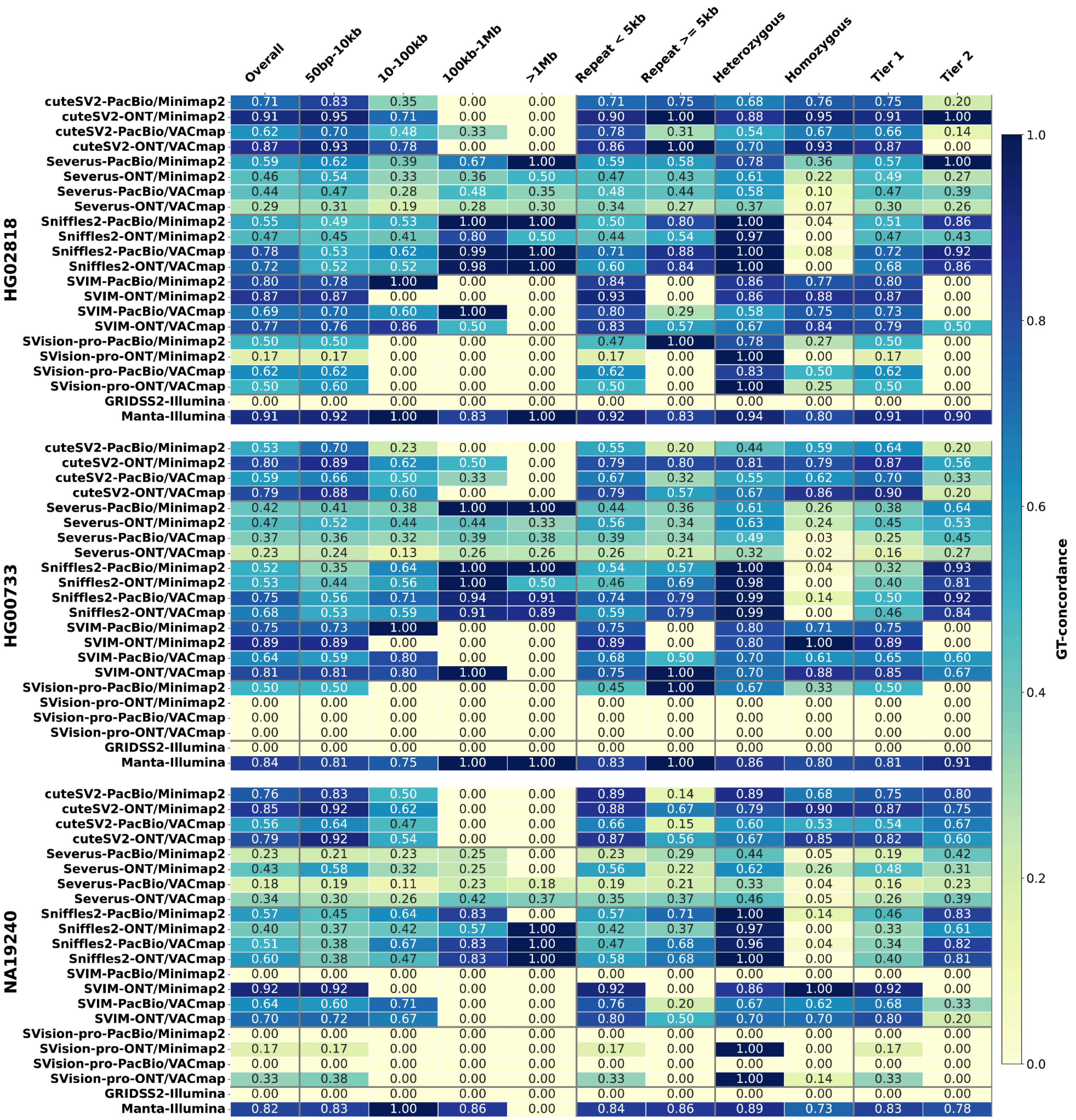


***Supplementary Figure 5: Genotype concordance of SV callers at 30x coverage of HG00733, HG02818, and NA19240 samples.*** *Each row represents a specific combination of a structural variant (SV) caller, sequencing technology (PacBio or ONT), and alignment software (Minimap2 or VACmap). Short-read callers using Illumina data are included below. The columns display GT-concordance scores stratified by genomic context, including overall performance, variant size, repeat element length, zygosity (heterozygous vs. homozygous), and evidence-based confidence (Tier 1 vs. Tier 2). The color intensity of each cell corresponds to the GT-concordance, with darker shades of blue indicating higher genotyping performance.*

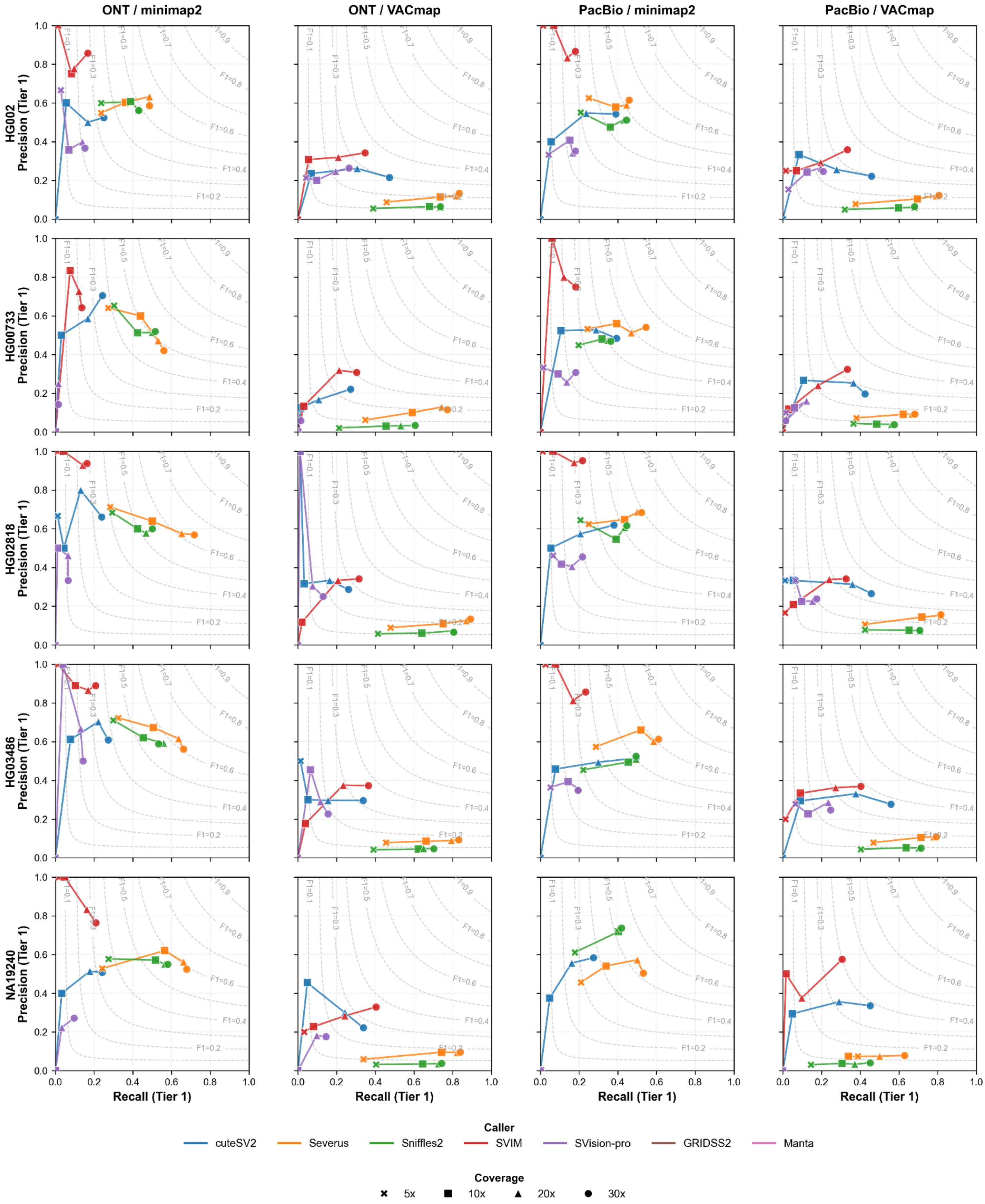


***Supplementary Figure 6: Subsampling effect on Tier 1 recall, precision, and F1 score on 30x data.*** *Each row represents a sample (HG002, HG00733, HG02818, HG03486, NA19240), each column represents a data*

*type (PacBio/minimap2, PacBio/VACmap, ONT/minimap2, ONT/VACmap). Each software is represented by one color to demonstrate the downsampling effect on precision, recall, and F1.*
